# Imaging-Genomics Study Of Head-Neck Squamous Cell Carcinoma: Associations Between Radiomic Phenotypes And Genomic Mechanisms Via Integration Of TCGA And TCIA

**DOI:** 10.1101/214312

**Authors:** Yitan Zhu, Abdallah S.R. Mohamed, Stephen Y Lai, Shengjie Yang, Aasheesh Kanwar, Lin Wei, Mona Kamal, Subhajit Sengupta, Hesham Elhalawani, Heath Skinner, Dennis S Mackin, Jay Shiao, Jay Messer, Andrew Wong, Yao Ding, Joy Zhang, Laurence Court, Yuan Ji, Clifton D Fuller, M.D. Anderson

## Abstract

**Purpose:** Recent data suggest that imaging radiomics features for a tumor could predict important genomic biomarkers. Understanding the relationship between radiomic and genomic features is important for basic cancer research and future patient care. For Head and Neck Squamous Cell Carcinoma (HNSCC), we perform a comprehensive study to discover the imaging-genomics associations and explore the potential of predicting tumor genomic alternations using radiomic features.

**Methods:** Our retrospective study integrates whole-genome multi-omics data from The Cancer Genome Atlas (TCGA) with matched computed tomography imaging data from The Cancer Imaging Archive (TCIA) for the same set of 126 HNSCC patients. Linear regression analysis and gene set enrichment analysis are used to identify statistically significant associations between radiomic imaging features and genomic features. Random forest classifier is used to predict two key HNSCC molecular biomarkers, the status of human papilloma virus (HPV) and disruptive TP53 mutation, based on radiomic features.

**Results:** Wide-spread and statistically significant associations are discovered between genomic features (including miRNA expressions, protein expressions, somatic mutations, and transcriptional activities, copy number variations, and promoter region DNA methylation changes of pathways) and radiomic features characterizing the size, shape, and texture of tumor. Prediction of HPV and TP53 mutation status using radiomic features achieves an area under the receiver operating characteristics curve (AUC) of 0.71 and 0.641, respectively.

**Conclusion:** Our analysis suggests that radiomic features are associated with genomic characteristics in HNSCC and provides justification for continued development of radiomics as biomarkers for relevant genomic alterations in HNSCC.

## INTRODUCTION

Head and neck squamous cell carcinomas (HNSCCs) prevail as the sixth most common cancer worldwide with over 500,000 expected newly diagnosed cases reported annually^1^. In the United States, 40,000 new HNSCC cases are reported with approximately 7,890 deaths per year^2^. HNSCCs encompass a diverse array of cancers that can originate from subsites within the oral cavity (44%), larynx (31%) or pharynx (25%)^3^. Viral infections, specifically human papilloma virus (HPV) primarily type 16 and Epstein-Barr virus, are associated with higher risk of oropharynx and nasopharynx cancers respectively^4-5^. Protracted tobacco and alcohol use as well as UV light exposure are among the traditional risk factors for development of HNSCC^6^. There has been a dramatic change in the affected patient cohort as risk factors has changed, represented by a decrease in tobacco use and concomitant increase in HPV-associated disease. This was reflected as a substantial rise in the incidence of HPV-associated oropharynx cancers as compared to a decline in cancers of the larynx and hypopharynx^7^. Given the high morbidity and mortality associated with HNSCC, this type of cancer represents a major health burden.

The refinement in head and neck irradiation techniques, specifically introduction of intensity-modulated radiotherapy about 15 years ago, was a paradigm shift HNSCC management that resulted in improvement of treatment outcomes^8^. Continued efforts have been made to investigate potential prognostic and predictive biomarkers to establish the conceptual framework for precision medicine in management of HNSCC^9^. One example is the exploration of the correlation between disruptive alteration of the gene encoding the tumor-suppressor protein p53 (TP53) and treatment failure with subsequent decreased survival in HNSCC patients^10^.

Radiographic images, such as Computed Tomography (CT), have been routinely used for diagnosis and treatment of HNSCC. However, the relationship between tumor imaging phenotypes and underlying tumor genomic mechanisms remains underexplored. Precise and effective treatment of cancer requires the integration of disease information from multiple sources. Imaging-genomics research combines radiographic image analysis with genomic research to improve disease diagnosis and prognosis, discover novel biomarkers, and identify genomic mechanisms associated with phenotype formation^11-15^. Such imaging-genomics studies have been performed for multiple cancer types, including breast invasive carcinoma^11-15^, lung cancer^16-17^, glioblastoma multiforme^18^, and clear cell renal cell carcinoma^19^.

To our knowledge, there are very few existing imaging-genomics studies for HNSCC. One of the earliest studies from 2003 by Yang et al. investigated the correlation between temporal changes in T1- and T2-weighted contrast-enhanced magnetic resonance imaging and genomic analysis using oligonucleotide microarrays in murine squamous cell carcinoma tumor models^20^. Aerts et al. developed a multi-feature radiomic signature capturing intratumoural heterogeneity that was linked to gene-expression patterns, validated in three independent data sets of lung and head-and-neck cancer patients^21^. Recently, Pickering et al. correlated radiologist-selected CT imaging features of 27 oral cavity squamous cell carcinomas with the expression of cyclin D1, angiogenesis-related genes, and epidermal growth factor receptors^22^.

In the current study, we innovatively investigated the comprehensive relationship between the multi-layer tumor genomic system and the multiple aspects of tumor imaging phenotype for HNSCC. We integrated multi-omics, whole-genome measurements from The Cancer Genome Atlas (TCGA)^23^ with radiomic data derived based on CT images from The Cancer Imaging Archive (TCIA)^24^ for matched patients, and identified statistically significant associations between them. We also explored the potential of using CT imaging as a non-invasive marker predicting the tumor molecular status for HNSCC.

## METHODS

Clinical, radiological, and genomic data (Supplemental Information Sections 1-2) for 126 HNSCC patients from TCGA and TCIA were integrated and analyzed. CT images of the patients were downloaded from TCIA and processed using Imaging Biomarker Explorer (IBEX)^25^, an automatic medical image analysis software pipeline that generates tumor radiomic features. The radiomic features were grouped into five categories: (1) gray level co-occurrence matrix, (2) gray level run length matrix, (3) neighbor intensity difference, (4) intensity direct, and (5) size/shape^21^. Supplementary Information Section 1 introduces how the radiomic features were generated. Multi-omics genomic data and patient clinical information were acquired from TCGA using the open-source R software tool TCGA-Assembler^26^. Supplementary Information Section 2 introduces the collection and processing of genomic data. Genomic data, clinical data, and radiomic data were integrated to form the imaging-genomics data (Table 1) for subsequent analysis.

**Table 1:**
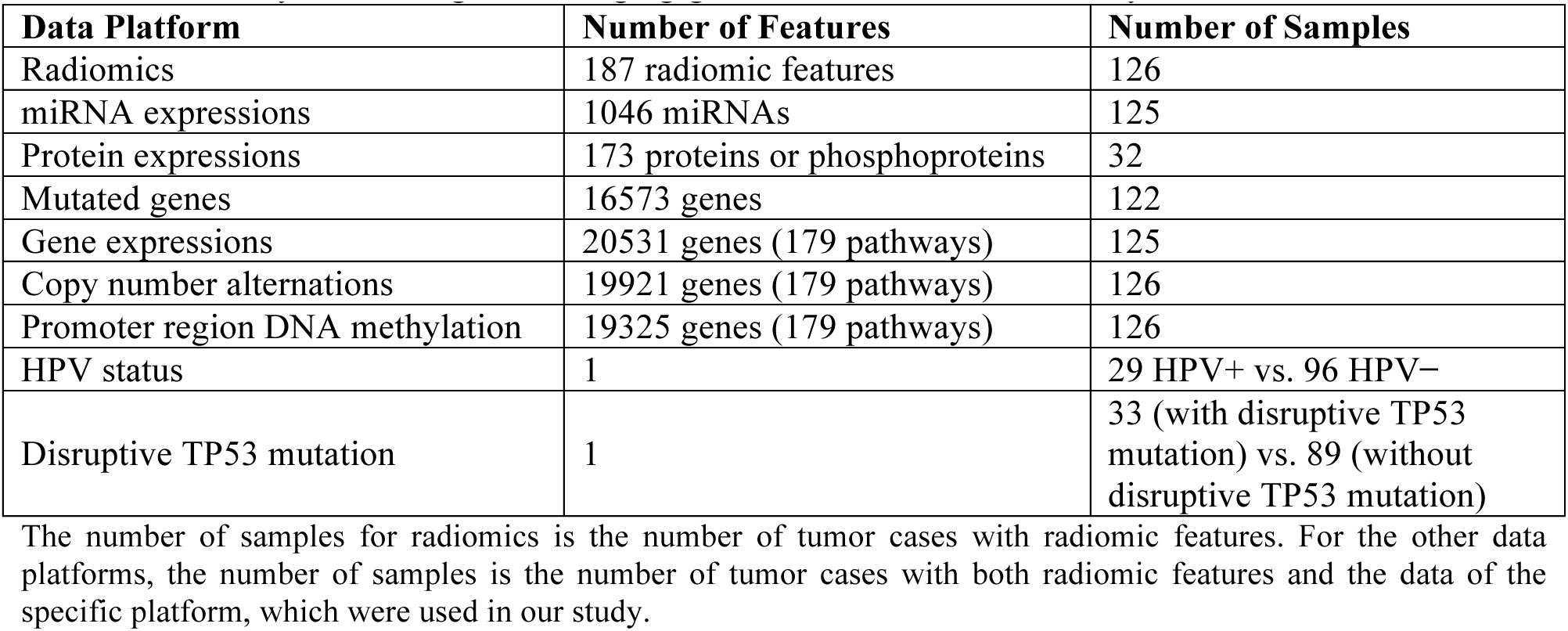
Summary of the integrative imaging-genomics data used in the analysis. The number of samples for radiomics is the number of tumor cases with radiomic features. For the other data platforms, the number of samples is the number of tumor cases with both radiomic features and the data of the specific platform, which were used in our study.

A multi-step informatic and statistical pipeline was built to perform integrative data processing and analysis (Fig. 1). First, linear regression was used to identify statistically significant associations between radiomic features and gene-level genomic features including expressions of miRNAs and proteins, and somatic mutations summarized at the gene level, adjusting for patient age, tumor grade, tumor subsite, and patient smoking status (Supplementary Information Sections 7-9). Second, for the whole-genome measurements, including gene expressions, copy number variations (CNVs), and promoter region DNA methylation, we investigated their associations with tumor radiomic features at the pathway level using a modified Gene Set Enrichment Analysis (GSEA)^27^ scheme that was also adjusted for the confounding factors mentioned above (Supplementary Information Sections 4-6). The genetic pathways in consideration are from the Kyoto Encyclopedia of Genes and Genomes (KEGG)^28^ database charactrerizing various aspects of the biomolecular system. Third, based on radiomic features, random forest classifers^29^ were used to predict patient HPV status and TP53 mutation status in HNSCC (Supplementary Information Section 10).

**Figure 1.**
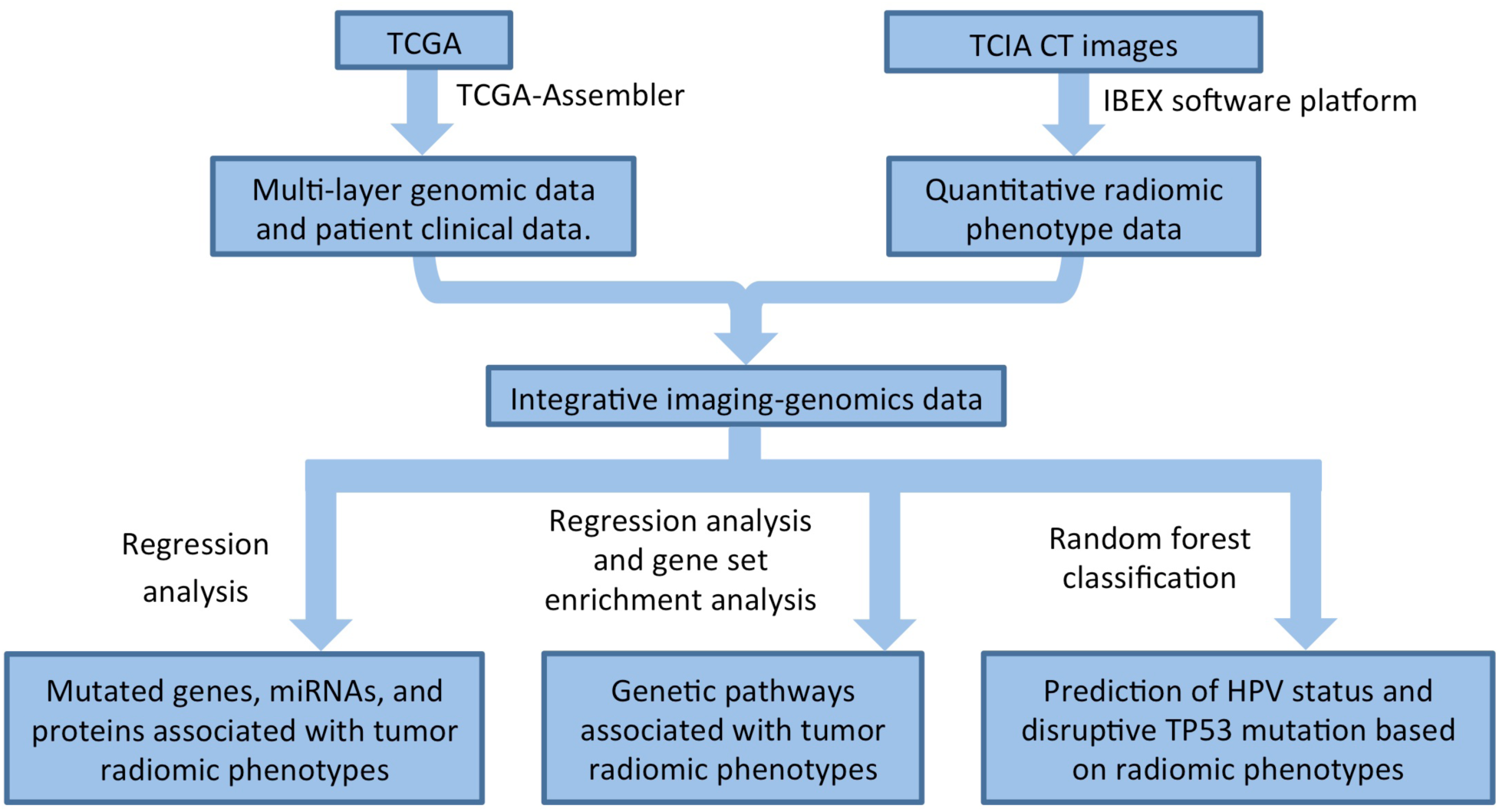
Flowchart of processing TCGA and TCIA data and conducting the imaging-genomics analyses.

## RESULTS

A total of 126 patient samples were analyzed, representing all matched cases in TCGA and TCIA HNSCC database(s), with AJCC stage IV (n = 86), stage III (n = 22), stage II (n = 14), and stage I (n = 4). The tumor subsites were oral cavity (n = 69), larynx (n = 36), and oropharynx (n = 21). Mean patient age was 59.81 years with a standard deviation of 11.28 years. Among all patients, 52 were current smokers, 45 former smokers, and 29 none smokers (never smoked). A total of 5,350 statistically significant associations (adjusted p-value ≤ 0.05) were identified between various radiomic and genomic features. Fig. 2a is a graphical presentation of the identified associations. Fig. 2b shows the numbers of identified associations between different categories of genomic features and radiomic features, based on which Fisher’s exact test^30-31^ indicates that the frequency of statistically significant associations depended on the feature category (p-value ≤ 1.0×10−^8^), meaning some feature categories have more associations than others. The identified associations are statistically significantly enriched among pathway transcriptional activities and all five categories of radiomic features with adjusted p-values < 1.0×10−^30^ (Table S3). This implies that transcriptional activities of genetic pathways modulate various aspects of tumor imaging phenotype.

**Figure 2.**
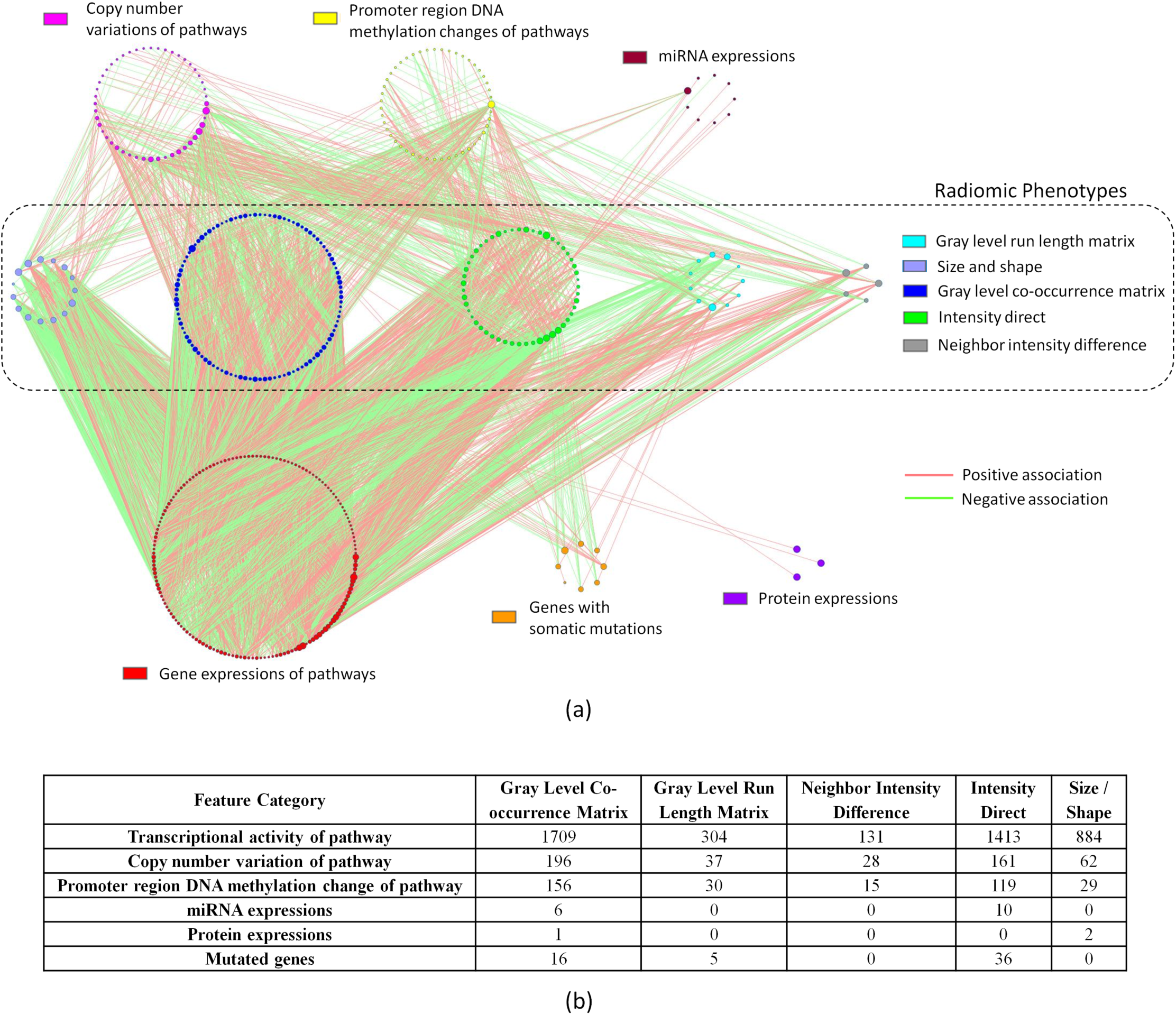
(a) Overview of all statistically significant associations identified in our analysis. Each node is a genomic or radiomic feature. Each line is an identified association. Genomic or radiomic features without significant association are not shown. Genomic features are organized into circles by data platform and indicated by different node colors. Radiomic features are divided into five categories also indicated by different node colors. The node size is proportional to its connectivity relatively to other nodes in the category. Associations are deemed as statistically significant if adjusted p-values ≤ 0.05. (b) Numbers of statistically significant associations between genomic features of different platforms and radiomic features of different categories.

### Associations between Radiomic Features and Genetic Pathways

Tables S4, S5, and S6 include all identified associations involving transcriptional activities, gene CNVs, and promoter region DNA methylation changes of all KEGG pathways, respectively. Fig. 3 specifically presents that radiomic features are associated with cancer-related KEGG pathways^28^ that cover multiple aspects of the cancer molecular system, such as signal transduction, cell growth and death, immune system, and cellular interactions and community. Fig. 3a, 3b, and 3c show the associations of transcriptional activities, gene CNVs, and promoter region DNA methylation changes of cancer-related KEGG pathways, respectively. There are many interesting findings in Fig. 3a indicating pathway transcriptional activities are correlated with and modulate multiple aspects of tumor imaging phenotype, and we elaborate on them below.

**Figure 3.**
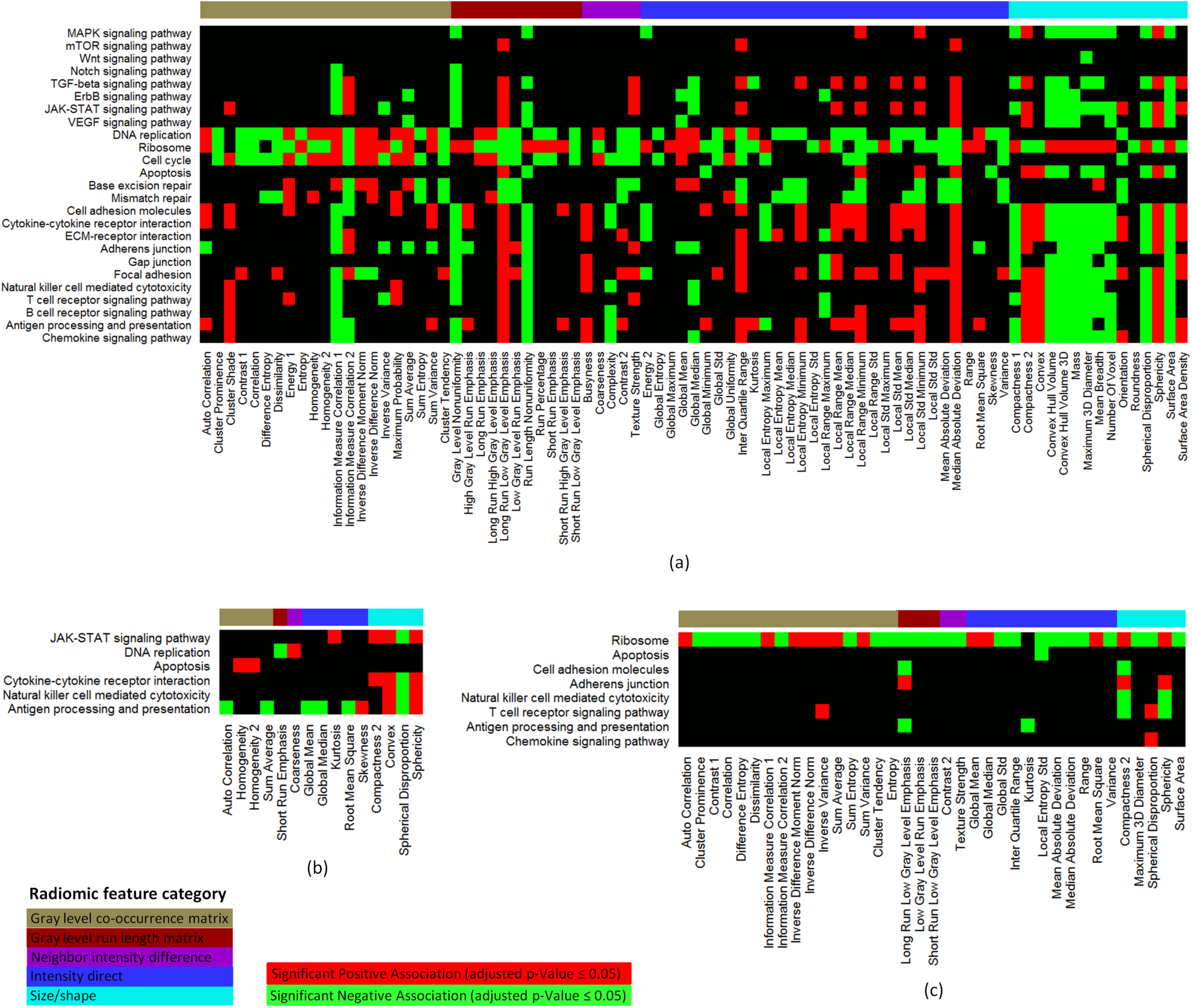
Statistically significant associations between radiomic features and (a) transcriptional activities of cancer-related genetic pathways, (b) gene CNVs of cancer-related genetic pathways, (c) gene promoter region DNA methylation changes of cancer-related genetic pathways. In each heatmap, only genetic pathways and radiomic features with statistically significant associations were shown. Each of the gray level co-occurrence matrix features can be calculated using different offset parameter values, i.e. 1, 2, 3, 4, and 5, which results in 5 different instances of a feature. Because the 5 instances of a feature were usually correlated, the directions (i.e. positive or negative) of the associations between a cancer-related pathway and the different instances of a radiomic feature were always the same. Thus, in the heatmaps, associations between different instances of a radiomic feature and a pathway could be collapsed into one association. If a pathway had an association with at least one instance of a radiomic feature, the association between the pathway and the radiomic feature was included in the heatmap. Percentile and quantile radiomic features from the intensity direct category were not included in the heatmaps for simplicity, because they have many instances with different percentile or quantile values.

### Cell Growth and Death

Multiple associations related to cell growth and death are identified in our analysis. Transcriptional activities of ribosome genes are correlated with multiple aspects of tumor imaging phenotype, including (1) tumor texture heterogeneity characterized by positive association with *entropy* and negative associations with *energy 1*, *homogeneity*, and *homogeneity 2*, (2) tumor size features, including *convex hull volume*, *convex hull volume 3D*, *mass*, *maximum 3D diameter*, *mean breadth*, *number of voxel*, and *surface area*, and (3) tumor shape irregularity, characterized by negative associations with *roundness*, *sphericity*, and *convex*, and positive association with *spherical disproportion*. Ribosome genes support protein synthesis and are important for various cellular processes, such as cell proliferation and growth. Our result shows that they are more transcriptionally active in larger, more irregular and heterogeneous tumors. The apoptosis pathway takes a tumor suppressive role by eliminating damaged or redundant cells through activating caspases. Disruption or evasion of apoptosis can lead to tumor initiation, progression or metastasis^32^. Consistently, we find that the transcriptional activity of apoptosis pathway is negatively associated with tumor size (characterized by *convex hull volume*, *convex hull volume 3D*, *maximum 3D diameter*, *mean breadth*, and *surface area*) and tumor shape irregularity (characterized by its positive associations with *convex* and *sphericity*, and negative association with *spherical disproportion*).

### Immune System

Pathways related to immune regulation, including pathways of natural killer cell mediated cytotoxicity, T cell receptor signaling, B cell receptor signaling, antigen processing and presentation, and chemokine signaling, are all negatively associated with tumor size features. One possible explanation is that patients with larger tumors have a less active immune system and therefore are unable to effectively destroy tumor cells and curb tumor growth. Similarly, we find a correlation between immune system activity and tumor shape regularity, as the pathway activities are positively associated with *sphericity* and *convex*, and negatively associated with *spherical disproportion*.

### Cellular Interactions and Community

Pathways related to cell adhesion molecules, cytokine-cytokine receptor interaction, ECM-receptor interaction, adherens junction, gap junction, and focal adhesion regulate cell-cell interaction and signaling acting as intercellular regulators and mobilizers of cells, and maintain cell and tissue architecture that limits cell movement and proliferation, which are two important factors in cancer progression. Aberrant activities of these pathways can lead to the development and metastasis of many types of cancer, including HNSCC^33^. We find that their activities are negatively associated with multiple tumor size features, indicating smaller tumors tend to have stronger activities of these pathways than large tumors. Activities of all these pathways, except gap junction, are also correlated with tumor shape regularity characterized by their positive associations with *sphericity* and negative associations with *spherical disproportion*.

### Signal Transduction

The transcriptional activities of several molecular signaling pathways, including MAPK signaling pathway, TGF-beta signaling pathway, JAK-STAT signaling pathway, VEGF signaling pathway, WNT signaling pathway, and ERBB signaling pathway, are negatively associated with tumor size features, indicating that they are more active in small tumors than large tumors. Previous report^34^ has suggested TGF-beta signaling as a potent tumor suppressor in HNSCC, which is supported by its negative association with tumor size identified in the current study. The activities of MAPK, TGF-beta, JAK-STAT, and VEGF signaling pathways are positively associated with tumor shape regularity.

Compared to pathway transcriptional activities, CNVs of cancer-related pathways have much fewer statistically significant associations with radiomic features (Fig. 3b). CNVs of JAK-STAT signaling pathway, cytokine-cytokine receptor interaction, natural killer cell mediated cytotoxicity, and antigen processing and presentation genes are correlated with tumor shape regularity characterized by their positive associations with *convex* and *sphericity,* and negative associations with *spherical disproportion*. CNVs of apoptosis genes are positively associated with tumor texture homogeneity characterized by *homogeneity* and *homogeneity 2*, indicating tumors with heterogeneous texture may have fewer copies of apoptosis genes than tumors with homogeneous texture.

Fig. 3c shows the statistically significant associations between radiomic features and promoter region DNA methylation changes of cancer-related pathways. DNA methylation changes of ribosome genes have the largest number of associations with radiomic features (first row in Fig. 3c), including negative associations with two tumor size features *maximum 3D diameter* and *surface area*, and positive associations with tumor shape regularity (characterized by positive association with *sphericity* and negative association with *spherical disproportion*). The directions of these associations are opposite of those for the transcriptional activities of ribosome genes, which is expected, since methylation at promoter region usually negatively affects gene expression. In addition, we find that DNA methylation changes of three immune related pathways, i.e. natural killer cell mediated cytotoxicity, T cell receptor signaling pathway, and chemokine signaling pathway, are negatively associated with tumor shape regularity (Fig. 3c). These are new results that may shed lights on the connection between immune pathways with radiomic phenotypes.

We report the analysis scheme and more findings in Supplementary Information Sections 4, 5, and 6.

### Associations between Radiomic Features and miRNA Expressions, Protein Expressions, and Mutated Genes

#### MiRNA

Table S7 presents statistically significant associations between miRNA expressions and radiomic features. *MiR-320a* has been reported as a negative regulator of tumor invasion and metastasis^35^. Its expression correlates with tumor texture homogeneity characterized by positive associations with *homogeneity* and *homogeneity 2* and negative associations with *entropy* and *global entropy*. The radiomic feature *global uniformity* measures the overall homogeneity of tumor pixel intensity^21^ and is positively associated with the expressions of 8 miRNAs including both antitumorigenic/antimetastatic and oncogenic miRNAs. The antitumorigenic/antimetastatic miRNAs include *miR-101* (targeting *EZH2*, a histone-lysine N-methyltransferase enzyme epigenetically silencing tumor suppressor genes^36^)*, miR-15b* (targeting *VEGF*, an important factor in the neo-angiogenesis process that is crucial for cells to reach and disseminate through the circulation system^37^), and *miR-320a*; the oncogenic miRNAs include *miR-106b* and *miR-25* (both from miR-106b-25 cluster that is over-expressed in HNSCC and promotes cell proliferation^38^), *miR-155* (upregulated in HNSCC and targeting tumor suppressors such as adenomatous polyposis coli^39^), and *miR-378* (reported to repress a potential tumor suppressor gene *TOB2* in nasopharyngeal carcinoma^40^); the last miRNA *miR-7* is involved in multiple cancer-related signaling pathways and has been reported with both oncogenic and antitumorigenic roles^38^.

#### Protein

TCGA provides the expression levels of 173 proteins or phosphoproteins, for which three statistically significant associations are identified (Table S8). ERK2 (encoded by *MAPK1*) is an important protein in the MAPK signaling pathway regulating cell proliferation, differentiation, and migration. Aberrant and/or persistent activation of the MAPK cascades can lead to the development and invasion of tumors including HNSCC^41-42^. The positive association between ERK2 expression and tumor *maximum 3D diameter* indicates larger tumors tend to have a higher ERK2 expression. The expression of Tuberin, a phosphorylation substrate of ERK2 encoded by *TSC2*, is also positively associated with *maximum 3D diameter.*

#### Somatic Mutation

Table S9 shows statistically significant associations between radiomic features and genes with somatic mutations in at least 10 patients. *EP300* encodes the E1A binding protein p300, a histone acetyltransferase regulating the transcription of genes involved in cell proliferation and differentiation. Mutations in *EP300* have been reported for HNSCC and may contribute to the disease initiation and progression^43^. Our analysis shows somatic mutations in *EP300* are negatively associated with *inverse variance* and positively associated with *median absolute deviation*. *COL11A1* encodes one of the two alpha chains of type XI collagen that is an essential component of the interstitial extracellular matrix. *COL11A1* may contribute to HNSCC tumorigenesis and be a potential therapeutic target^44^. We find mutations in *COL11A1* are negatively associated with *inverse variance*.

We report the analysis scheme and more findings in Supplementary Information Sections 7, 8, and 9.

### Predictions of Patient HPV Status and Disruptive TP53 Mutation Using Radiomic Features

We applied the random forest classifier^29^ to predict the patient HPV status based on tumor radiomic features. A two-tier five-fold cross-validation was used to tune the classifier parameters and evaluate the generalization prediction performance. Predictive radiomic features were selected through a recursive feature elimination scheme. Table 2 shows the mean and standard deviation of the Area Under the receiver operating characteristic Curve (AUC) across 30 cross-validation trials, which measures the prediction accuracy. There is no significant difference between the average AUCs obtained using different numbers of features for prediction. The highest average AUC achieved is 0.71, while the average AUC using only five features in each cross-validation trial still reaches 0.706. Using the same classification and feature selection scheme, we predicted whether a tumor possessed any disruptive TP53 mutation, a biomarker in HNSCC development and treatment^10^. Table 2 shows the mean and standard deviation of obtained AUCs. The highest average AUC is 0.641 with five features selected for prediction in each cross-validation trial. See the Supplementary Information Section 10 for details of the prediction and feature selection scheme, and additional details of results.

**Table 2:**
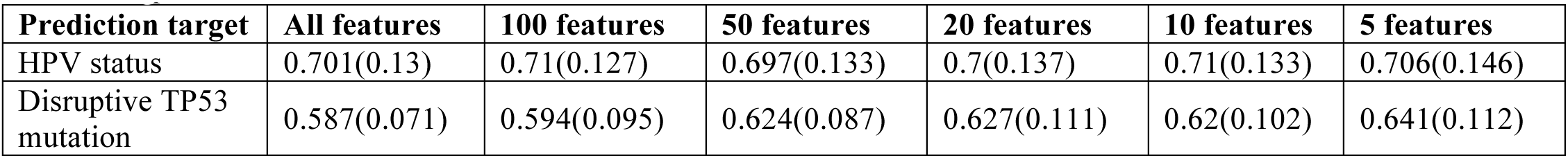
Mean (standard deviation) of AUCs obtained through a two-tier five-fold cross-validation scheme that includes 30 cross-validation trials when different numbers of radiomic features were selected for prediction in each cross-validation trial.

## DISCUSSION

Using TCGA and TCIA data, we conducted a comprehensive imaging-genomics study. To our knowledge, this is the first study that integrates radiomic features of CT images with whole-genome measurements depicting multiple layers of tumor molecular system for HNSCC. We report statistically significant associations between radiomic features characterizing multiple aspects of the tumor imaging phenotype and various genomic features (including transcriptional activity, CNV, DNA methylation, miRNA expression, protein expression, and somatic mutation). The identified associations support existing knowledge related to HNSCC pathogenetic mechanisms and provide evidence for novel hypotheses on the potential relationship between tumor genomic mechanisms and subsequent tumor phenotypes. Also, we attempted to use radiomic features to predict important molecular biomarkers in HNSCC, such as HPV status and disruptive TP53 mutation, with decent AUC values. These results provide basis for future investigations to establish the potential of using non-invasive imaging approach to probe the genomic and molecular status of HNSCC. Our findings are uploaded to http://www.compgenome.org/Radiogenomics/ as a public resource to facilitate future research on HNSCC imaging-genomics.

Compared to pathway transcriptional activities, much fewer statistically significant associations have been identified for pathway CNVs and DNA methylation changes (Fig. 2b and Fig. 3). There could be two reasons for this. First, transcriptional activity is closer to phenotype formation than CNV and DNA methylation in the process of molecular system regulating the development of phenotype. Basically, transcriptional activities can more directly influence the generation of various phenotypes, while CNVs and DNA methylation changes may have to function through transcription. Secondly, DNA mutation events, such as CNVs and somatic mutations, are rarely shared across many patients, resulting in a small number of samples with the same mutation event that limits the statistical power for identifying potential associations.

Our study is based on CT images of 126 HNSCCs and their multi-layer whole-genome genomic data, which form a unique imaging-genomics dataset that was not available before TCGA/TCIA era. Although this dataset is so far the largest of its kind, its sample size might still limit the statistical power for identifying imaging-genomics associations and the accuracy of predicting tumor molecular status based on radiomic features. Nonetheless, we believe our study will pave ways for future HNSCC imaging-genomics investigation using more samples and more imaging technologies.

More imaging-genomics analyses have been planned for HNSCC. One particularly interesting approach is to integrate genomics, epigenomics, and proteomics data simultaneously with imaging data to provide a more comprehensive depiction of how the multi-layer molecular system regulates and produces various tumor imaging phenotypes. Graphical models can be powerful tools for studying such complex relationship, due to their ability to model conditional dependence and competing regulatory factors^45^.

## Authors’ Contributions

All listed co-authors performed the following. 1. Contributions to the conception or design of the work; or the acquisition, analysis, or interpretation of data for the work. 2. Drafting the work or revising it critically for important intellectual content. 3. Final approval of the version to be published. 4. Agreement to be accountable for all aspects of the work in ensuring that questions related to the accuracy or integrity of any part of the work are appropriately investigated and resolved. Specific individual cooperative contributions to study/manuscript design/execution/interpretation, in addition to all criteria above are listed as follows. YZ: manuscript writing, acquisition and preprocessing of The Cancer Genome Atlas (TCGA) data, integration of TCGA data and radiomics data, conceived and conducted all statistical analyses, interpretation of analysis results. ASRM: manuscript editing, direct oversight of image segmentation and image post-processing, and clinical data collection workflows; direct oversight of trainee personnel. SY: construction of website resource hosting the identified imaging-genomics associations. HE, MK: manuscript editing, oversight of imaging/clinical data collection overflows. LW: implementation of software pipeline for acquisition and preprocessing of TCGA data. AK, DM, JZ, LC: development support for radiomics workflow and curation of the radiomics-based image features and relevant clinical data. SS: participation in statistical analysis of imaging-genomics data. DM, JM, AW, YD, AK: TCIA/TCGA records screening, automated case identification, data extraction, imaging/clinical data collection and informatics software support. AK, JS, LC, HE: clinical data curation, image segmentation, data transfer and supervision of DICOM-RT analytic workflows. SYL, HS: database curation and oversight, supervisory support, editorial oversight; genomics conceptual feedback and support. CDF, YJ: corresponding author; primary investigator; conceived, coordinated, and directed all study activities, responsible for data collection, project integrity, manuscript content and editorial oversight and correspondence; direct oversight of trainee personnel.

